# MRI and cognitive scores complement each other to accurately predict Alzheimer’s dementia 2 to 7 years before clinical onset

**DOI:** 10.1101/567867

**Authors:** Azar Zandifar, Vladimir S. Fonov, Simon Ducharme, Sylvie Belleville, D. Louis Collins, for the Alzheimer’s Disease Neuroimaging Initiative

**Author notes:** **Corresponding Author Information:** Azar Zandifar, McConnell Brain Imaging Centre, Montreal Neurological Institute Montreal Neurological Institute, 3801 University Street, Room WB320, Montreal, QC, H3A 2B4.

## Abstract

**Background:** Predicting cognitive decline and the eventual onset of dementia in patients with Mild Cognitive Impairment (MCI) is of high value for patient management and potential cohort enrichment in pharmaceutical trials. We used cognitive scores and MRI biomarkers from a single baseline visit to predict the onset of dementia in an MCI population over a nine-year follow-up period.

**Method:** All MCI subjects from ADNI1, ADNI2, and ADNI-GO with available baseline cognitive scores and T1w MRI were included in the study (n=756). We built a Naïve Bayes classifier for every year over a 9-year follow-up period and tested each one with Leave one out cross validation.

**Results:** We reached 87% prediction accuracy at five years follow-up with an AUC>0.85 from two to seven years (peaking at 0.92 at five years). Both cognitive test scores and MR biomarkers were needed to make the prognostic models highly sensitive and specific, especially for longer follow-ups. MRI features are more sensitive, while cognitive features bring specificity to the prediction.

**Conclusion:** Combining cognitive scores and MR biomarkers yield accurate prediction years before onset of dementia. Such a tool may be helpful in selecting patients that would most benefit from lifestyle changes, and eventually early treatments that would slow cognitive decline and delay the onset of dementia.

## Introduction

Alzheimer’s disease (AD) is a progressive neurodegenerative disease [1–3]. The prodromal stage of AD dementia, known as Mild Cognitive Impairment (MCI), is characterized by the gradual onset and evolution of cognitive impairment beyond the levels expected for age and education of the individual, but without interfering with a patient’s everyday life [4]. MCI patients with memory problems as their main symptom are known as “amnestic MCI”, from which 10%-15% are reported to progress to clinically probable AD each year [5]. Since not all amnestic MCI patients progress to AD, predicting if and when a subject with MCI will have future dementia will enable enrichment for clinical trials. More importantly, Kivipelto’s group [6] has demonstrated the potential benefit of combining diet, exercise and cognitive training to prevent cognitive decline. Early detection of prodromal disease (e.g., 5-10y in advance) is key to intervene before the onset of cognitive decline due to irreversible neurodegeneration.

Jack *et al’s* [8] well-known hypothetical biomarker model and its successors [7] describe the dynamics of biomarkers during AD pathological process. The model proposes that cognition and structural brain atrophy change with the sharpest slope in the MCI stage [7], making them potentially the most sensitive early biomarkers of progression from MCI to AD. According to this model, anatomical atrophy, which can be captured by volumetric MRI, begins ahead of cognitive decline [7]. Specifically, the entorhinal and hippocampal areas are known to be among the first affected [9, 10]. Factoring the sharp slope of atrophy, preclinical volume loss and its widespread availability, MRI morphometric analyses is a promising candidate biomarker for earlier prediction.

To measure AD-related morphological changes in brain anatomy, we have developed the Scores by Nonlocal Image Patch Estimator (SNIPE) for both hippocampal and entorhinal areas [11, 12]. SNIPE is a similarity metric that measures structural similarity to either a library of cognitively normal subjects or a library of patients with Alzheimer’s dementia. We previously showed that using only SNIPE scores for hippocampus, plus age and sex, one can reach an overall accuracy of 71% for prediction of progression from MCI to dementia over a 3y follow-up period [11]. In a more recent work looking at a cohort of cognitively healthy older individuals, we showed that our MR-driven SNIPE biomarker was sensitive to AD-related changes in a cognitively normal cohort on average seven years before clinical diagnosis of AD dementia [13].

While volumetric MRI measures have good prediction value, we hypothesize that prediction performance can be improved using complementary information of other features, in particular performance on standardized cognitive tests. Indeed, previous studies showed that the patient’s current cognitive state can also predict future cognitive decline in MCI [14, 15]. The ACE-R (Addenbrooke’s Cognitive Examination - Revised) [16], MOCA (Montreal Cognitive Assessment) [17], verbal memory measures and many language tests [14] all have shown promising performance to predict future dementia in persons with MCI. However, some models may have difficulty with short term prediction as they found the likelihood of false negatives was increased, resulting in decreased sensitivity to imminent onset of AD [14]. Furthermore, these studies either suffer from relatively short follow-up periods, or have not used a combination of different scores and biomarkers to benefit from their complementary information. While a more recent study shows that combining both cortical thinning measures and cognitive scores increases prediction accuracy [18], this study too had a relatively short follow-up period, and a limited number of subjects and did not investigate the complementary effect of the different features [18].

While both SNIPE and cognition have shown potential to predict conversion, it is uncertain if a combination of both measures would lead to superior accuracy. In this study we have two main goals. First, we investigate the ability of our model to predict progression to AD based only on the baseline MR and cognitive information over follow-up periods ranging from one to nine years. Second, we investigate the change over time in predictive power of each feature to investigate whether, as previously suggested in Jack’s model [8], the MR-driven biomarkers are better predictors than cognitive tests for longer follow-up periods (as atrophy is hypothesized to precede cognitive decline), and whether they can capture AD-related abnormality before cognitive scores do. This study further investigates the effect of combining neurocognitive scores and MRI makers throughout different follow-up periods using a large dataset.

## Methods

In this study, we train a Naïve Bayes classifier to predict the future diagnosis of dementia in patients with MCI in the ADNI1, ADNI2 and ADNI-GO datasets. Our feature set contains age, sex, cognitive (Alzheimer’s Disease Assessment Scale (ADAS), Rey Auditory Verbal Learning Task (RAVLT), Mini-Mental State Examination (MMSE)), MR-based scores (SNIPE scores for hippocampus and entorhinal cortex), and disease severity (CDR-SB (Clinical Dementia Rating Sum of Boxes)) from baseline data that are used as input to the classifier. The classifier then attempts to predict the future diagnosis for each patient. To train the classifier, we need patient diagnostic information collected at later time points during the study. The follow-up period is the time interval from baseline to later time points for which we know the state of the subject: i.e., whether that subject has progressed to dementia or has maintained the MCI stage.

### Dataset selection

Data used in this study were obtained from the Alzheimer’s Disease Neuroimaging Initiative (ADNI) database (adni.loni.usc.edu). The ADNI was launched in 2003 as a public-private partnership, led by Principal Investigator Michael W. Weiner, MD. The primary goal of ADNI has been to test whether serial magnetic resonance imaging (MRI), positron emission tomography (PET), other biological markers, and clinical and neuropsychological assessment can be combined to measure the progression of mild cognitive impairment (MCI) and early Alzheimer’s disease (AD).

We used all MCI subjects from ADNI1, ADNI2, and ADNI-GO for which the baseline cognitive scores and T1 MRI were present (n=756). We used MRI and cognitive data from the baseline visit to predict the future clinical diagnostic status at follow-up at 12, 24, 36, 48, 60, 72, 84, and 108 months (See Table 1). (While 120-month data is available, too small a number of subjects are available for model testing and validation.)

**Table 1.** Dataset Information based on follow-up duration. sMCI stable MCI subjects, while progressive MCI population are referred to as pMCI. All information reported is based on baseline visit for each follow-up cohort.

|  | 12<br>months | 24<br>months | 36<br>months | 48<br>months | 60<br>months | 72<br>months | 84<br>months | 96<br>months | 108<br>months |
| --- | --- | --- | --- | --- | --- | --- | --- | --- | --- |
| **pMCI**<br>**subjects (#)** | 101 | 174 | 174 | 127 | 70 | 53 | 32 | 23 | 17 |
| **sMCI**<br>**subjects (#)** | 619 | 431 | 325 | 200 | 97 | 50 | 44 | 33 | 21 |
| **pMCI:sMCI**<br>**ratio** | 0.163 | 0.404 | 0.535 | 0.635 | 0.722 | 1.060 | 0.727 | 0.697 | 0.810 |
| **Mean Age**<br>**at baseline** | 73 | 73 | 73 | 72 | 72 | 74 | 74 | 74 | 73 |
| **Female**<br>**Percentage**<br>**(%)** | 39 | 40 | 38 | 38 | 35 | 35 | 34 | 29 | 34 |
| **Median**<br>**education** | 16 | 16 | 16 | 16 | 16 | 16 | 16 | 16 | 16 |

At each visit of the ADNI study, patients are evaluated for AD based on NINCDS-ADRDA criteria [3]. MCI participants are identified as those that have reported a subjective memory concern either autonomously or via an informant or clinician; have abnormal performance on the Wechsler Memory Scale Logical Memory II test, however activities of daily living are preserved (CDR=0.5) and they do not meet criteria for a dementia diagnosis.

The subjects were labeled either stable MCI (sMCI) or progressive MCI (pMCI) label based on the difference in diagnostic status between the baseline visit and each follow-up time point. For example, at 24 months, we included all MCI subjects for which the diagnostic state at 24 months was present. The subjects who maintained their baseline MCI state were labeled sMCI. Subjects that progressed to an AD diagnosis at any time *up to and including* the follow-up time point received the pMCI label. (Note: we do not consider later time points, as our goal is to *match the clinical status at the given time point*.) When a subject did not have a diagnostic label for a time point, that subject was dropped from the analysis for that follow-up time point only. The detailed MCI dataset information is given in Table 1. The strategy used here to categorize the data prevents the bias that could have been introduced to the analyses given the clinical status of the subjects who dropped out from the study.

### Feature set

#### MR derived biomarker

Hippocampal and entorhinal SNIPE grading scores were used as the only MRI biomarker features in the predictive classifier. The SNIPE score is described fully in [13, 20]. In short, SNIPE assigns a similarity metric to each voxel, which shows how much that voxel’s neighbourhood resembles the anatomy of either a group of patients with Alzheimer’s dementia or a group of normal controls. The final SNIPE score is an average of all the voxels in the desired anatomical structure in each hemisphere [12, 13]. For this study the SNIPE scores are corrected for age and sex using the method presented in [21] based only on the cognitively normal population to correct for the effect of normal aging and preserve the effect of disease-related changes. All processed MRI was submitted to visual quality control. No datasets needed to be excluded for insufficient quality.

#### Neurocognitive Scores

Previous studies showed that the prognostic value of cognitive tests can vary depending on remaining time to future onset of dementia [14]. Here, we included the baseline cognitive test scores available within the ADNI study. These included the total score of ADAS (both ADAS11 and ADAS13), the Rey Auditory Verbal Learning Task (RAVLT) scores of immediate, learning and forgetting, and MMSE. We also included baseline CDR-SB. The values are corrected for age, sex and education using the method presented in [21].

#### Classification and validation

We trained three classifiers based on the following features: first, using only four MR-driven biomarkers (both left and right entorhinal and hippocampal SNIPE score) and age and sex information; second, using only seven neurocognitive scores plus age and sex; and third, using all MR driven and cognitive scores together plus age and gender. All features are drawn only from the baseline visit. For each follow-up, we trained a specific Naïve Bayes classifier, for which the baseline information is used as features and the stable/progressing state of a patient at that specific follow-up period is regarded as the desired output. All classifiers are tested with a Leave One Out (LOO) cross-validation technique. This means that in each step, the classifier is trained on all the subjects from the dataset for that specific follow-up except one specific subject that is used as the test subject. This process is repeated so that all subjects are used for testing. Therefore, we have a completely separate testing and training sets.

### Metric

#### Classification accuracy, sensitivity, specificity and Area under the Receiver Operating Curve (AUC)

The classification performance at each time point is measured by classification accuracy, sensitivity and specificity for each follow-up period. To make the comparison between the metrics for different settings (using only MR-driven scores, only using cognitive scores, or using both sets of features) feasible and to gain a more robust sense of the classification performance, we measured the classification accuracy, sensitivity and specificity by sampling 85 percent of data without replacement for each iteration and we repeated the performance calculation procedure for 200 iterations. That is, for each iteration, we ran a LOO procedure only using 85% of the population. The final accuracy, sensitivity and specificity shown are the mean of the calculated metrics and the standard deviations is shown with error-bars.

As a measure that is robust to unbalanced classes, we also report the AUC for the classifier trained with all the features to better show our model performance.

#### Feature importance

To determine the relative importance of the different features over the 9-year follow-up period, we use a metric similar to effect size, customized for Naïve Bayes classification. In summary, we compare the different features based on their importance for our classifier at each point over time in the follow-up period (see the supplementary material for details). This shows how the features gain or lose importance over time and is related to the disease progression.

## Results

### Classification Performance

The accuracy, sensitivity and specificity at each follow-up time point are plotted in Fig. 1 for the three classifier scenarios: using only MRI features, using only cognitive scores and using both MRI and cognitive features together as inputs. From 24 months onwards, the classification accuracy using both sets of features are better than using only MR features or only cognitive features. MRI features are more sensitive (Fig. 1, middle), while cognitive features bring specificity to the prediction (Fig. 1, right).

**Figure 1.**
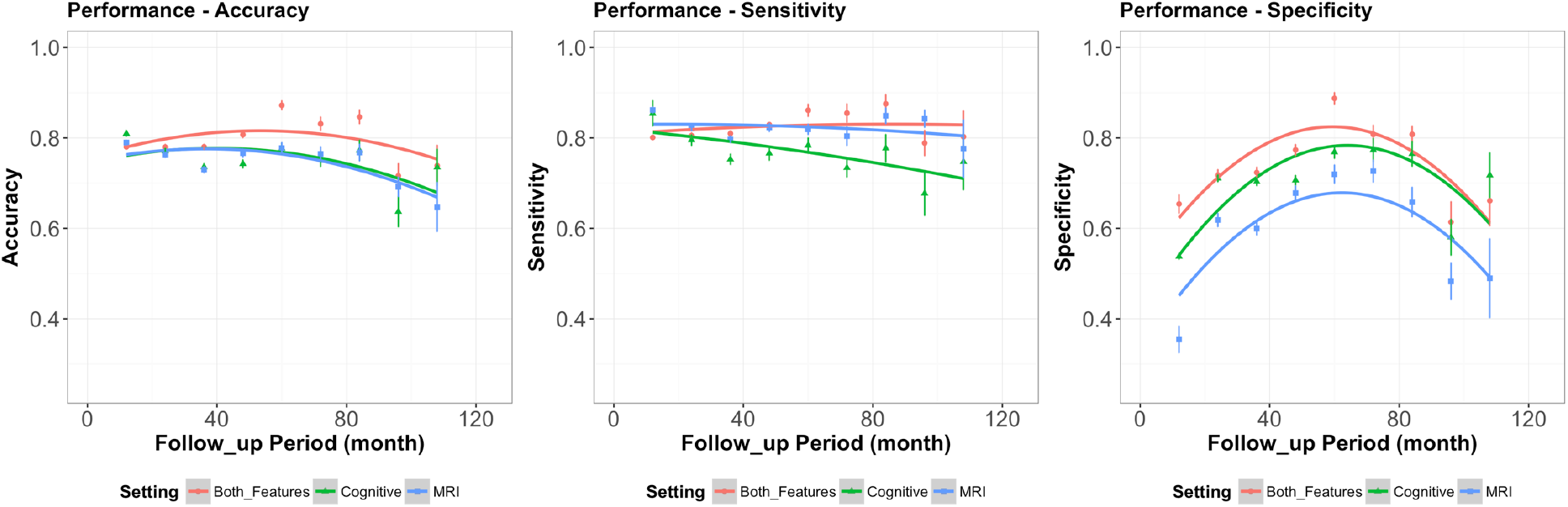
Classifier performance -- From left to right, Accuracy, Sensitivity and Specificity for follow-ups from 12 to 108 months. Data points show the mean metric calculated in each follow-up period from 200 bootstrap samples of 85% of the data. Error bars show the associated standard deviation. The curves show the best second-degree fit.

Statistical comparisons were made between accuracies, sensitivities and specificities of all the classifiers with different feature sets. Models were compared when trained using combined feature sets (i.e. MRI and cognitive scores) and with each one of them alone. An ANOVA with post-hoc pairwise Tukey test with 95% confidence level shows that the combined model significantly performs better than the models trained on each feature set alone in all follow-up periods (*p*<2× 10^−5^) except for the accuracies of the combined model and the model trained with only cognitive scores at 108 months.

As stated in the methods section, data imbalance may cause bias towards one of the classes, thus affecting the classification results. For example, at 12mo, there are only 101 pMCI and 619 sMCI. A simple classifier that assigns all the test cases to the majority class would obtain 86% accuracy by assigning all subjects to sMCI class. We therefore measured AUC, which is a more robust metric against data imbalance (Fig. 2). The AUC plot over time shows an inverted-U pattern where the maximum performance happens around 60 months (*AUC* = 0.92) with accuracy, sensitivity and specificity of 87%, 86%, and 89%, respectively. The model is robust with AUC values >0.85 for follow-up periods from 24 to 84 months.

**Figure 2.**
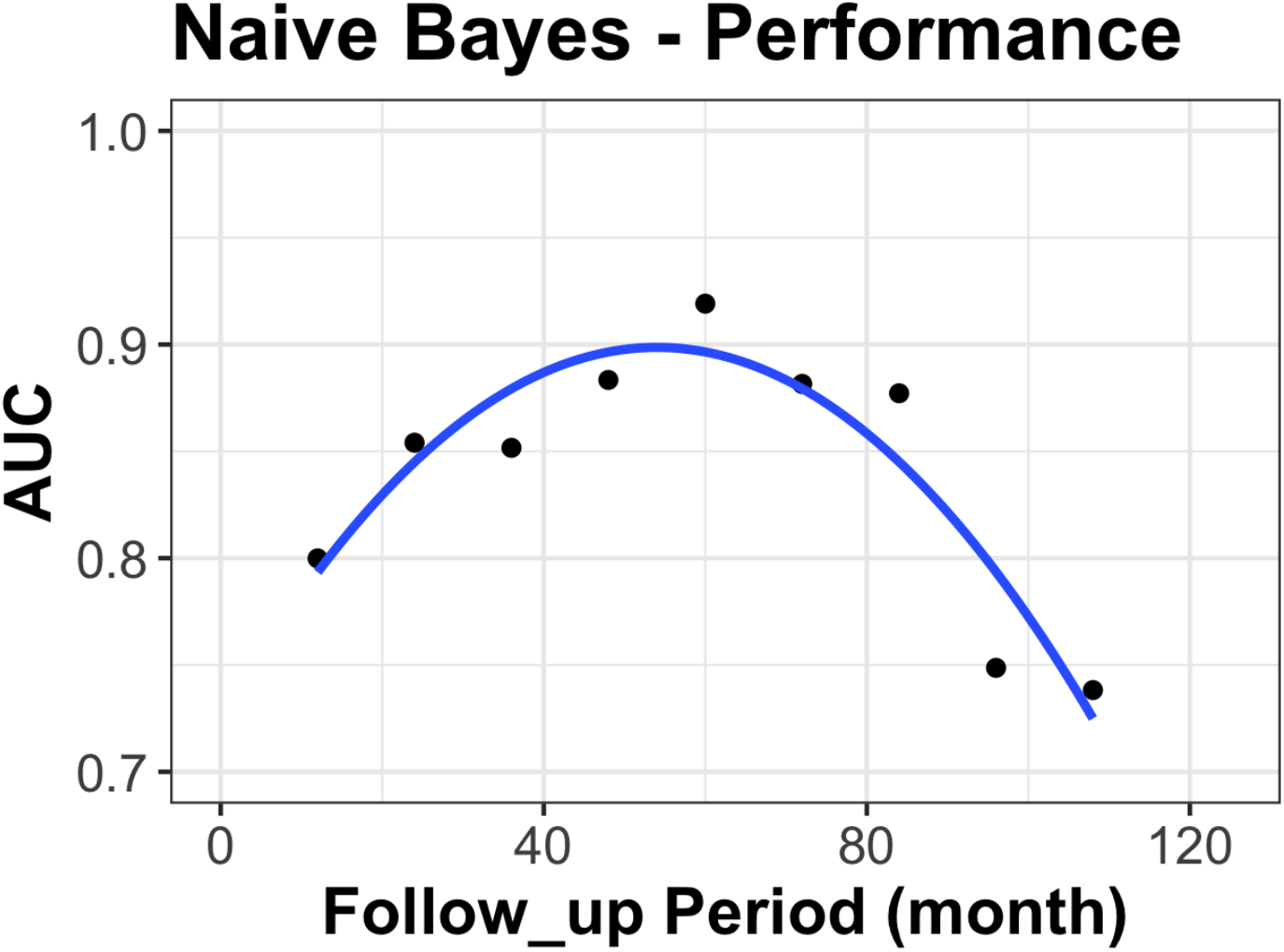
Area under the receiver operating curve (AUC) for follow-ups from 12 to 108 months. Data points show the metric in each follow-up period. The curve shows the best second-degree fit (Akaike Information Criterion yielded -38.12 for second order vs. -19.95 for a linear model).

The inverted-U pattern in the AUC plot (Fig. 2) is counterintuitive as baseline visits should be more informative for early follow-up periods and less informative for longer follow-up periods and, therefore, model performance should fall monotonically in time. Indeed, the sensitivity profile drops over time for the classification models using only MR or only cognition features but remains relatively flat for the combined model. The inverted-U pattern of the AUC curve (Fig. 2) is driven by the inverted-U shape of the specificity plot (Fig. 1), indicating that the classifier does not properly identify the negatives (sMCI) at the earlier follow-up time points. Early follow-ups are prone to more false positives. We hypothesize that this happens due to the fact that shorter follow-ups don’t allow enough time for all prodromal AD subjects to progress to a clinical dementia diagnosis. This means that these false positives are individuals who do not change categorical diagnostic categories in shorter follow-ups even though the disease is progressing and will reach the dementia stage at a later time point. To test this hypothesis, one could show that the conversion rate in this group is greater than the average MCI population. From the 110 false positives at 12 months, 71 (64%) converted to dementia, with more than 42% converting within one year (see Table 2 for details). This is much higher than the expected yearly rate of progression of 10-15% [5], indicating that many of these false-positives are in fact true progressors at a later follow-up time.

**Table 2.** False positive at short follow-up investigation through future follow-ups. FP stands for False Positives

|  | Total of<br>FP<br>(#) | Total Late<br>convertors<br>(#) | Convertors at<br>24 months<br>(#) | Convertors at<br>36 months<br>(#) | Convertors at<br>48 months<br>(#) | Convertors<br>after 60<br>months<br>(#) |
| --- | --- | --- | --- | --- | --- | --- |
| **12**<br>**months** | 110 | 71 | 47 | 12 | 8 | 4 |

### Feature importance

Importance of each feature in predicting disease progression at each follow-up is shown in Figure 3. For illustration, we averaged importance over ADAS 11 and 13, RAVLT learning, forgetting and immediate, hippocampal SNIPE score for left and right, entorhinal SNIPE score for left and right, and made one score out of each set. This graph shows the value of each feature in making the final decision when the classifier has access to both cognitive and MR data. ADAS shows the greatest importance over the entire follow-up period, followed by hippocampal SNIPE scores.

**Figure 3.**
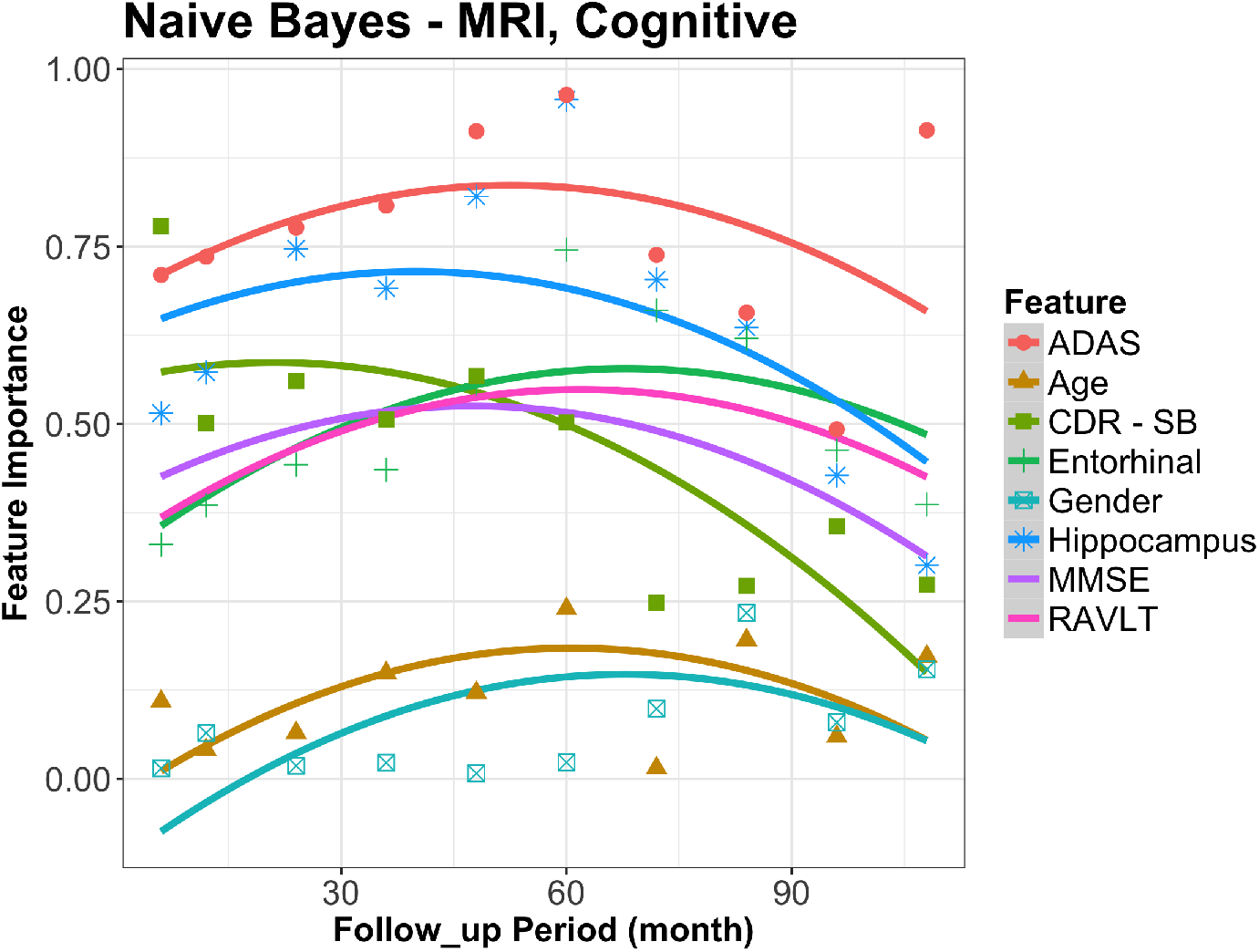
Feature importance for each follow-up period. Features show both cognitive and MR driven scores. The curves show the best second-degree fit given the metric for the 9-year follow-ups.

While all features lose AD-related sensitivity for very long follow-ups, the peak importance for each one happens at a different follow-up period. The later the peak, the more sensitive a feature would be to earlier detection of the disease. If used alone, entorhinal cortex shows the earliest possible detection performance at 68 months and is among the first biomarkers to show abnormality. While the hippocampal grading score has a peak at 40 months, it shows a higher importance for all the follow-up periods from one to seven years compared to the entorhinal cortex SNIPE score. Similarly, by peaking at 62 months, the RAVLT cognitive scores are more sensitive to early AD changes in comparison to the more general screening test MMSE and the disease severity marker CDR-SB (performance peak at 47 and 21 months, respectively). Having multiple biomarkers available enables each classifier to select the most appropriate data to achieve high sensitivity and specificity for all follow-up periods.

The demographic information shows very low importance in comparison to all the other features (Figure 3). In fact, their effect is negligible with respect to the other markers. However, they show higher importance for late follow-ups rather than short ones (peaks at 61 months and 67 months for age and sex, respectively).

## Discussion

In this study, we evaluated a prognostic model to predict the onset of dementia using baseline cognitive and MR-driven features from ADNI subjects as a function of follow-up period.

This study showed that both MR and cognitive information are needed to make the most accurate decision on future progressors. Importantly, they contribute differently to overall accuracy. The MR-driven features are more sensitive than cognitive scores, but cognitive scores are more specific than MRI features. In other words, the MR derived biomarkers are highly sensitive to the morphological pattern of abnormal brain aging (i.e., to identify at-risk subjects), while preservation of cognition function contributes to identify persons who will not progress over time despite MRI changes. This would explain why using both make an optimally accurate, sensitive, and specific prognostic tool.

One of the observations in this study was that the AUC follows an inverted-U shape over the 9-year follow-up period: at the beginning, AUC rises with time, reaches a maximum, and then falls again. It has been previously observed that a model trained based on cognitive scores can show low predictive ability for shorter follow-up periods [14]. This study reported low sensitivity in shorter follow-ups using some cognitive scores such as semantic fluency tasks [14]. By combining many of the cognitive scores available in ADNI with MRI features we maintained high sensitivity throughout the 9-year follow-up period. The study postulates that in shorter follow-ups, performance is adversely affected by late converters (i.e. individuals whose progression is not sufficient at the one-year time point to change diagnostic category but reach the dementia stage later). However, to the best of our knowledge, no previous study has proven the late-converters hypothesis and their effect on model performance using a data-driven approach. Our investigation with short follow-ups showed that the majority of false positives are in fact late convertors. Indeed, 42% of the false positives at 12 months convert to dementia within the following year. This is almost three times the rate of conversion to dementia found in the average MCI population [5]. More specifically, when our model falsely predicts the conversion to dementia at 12 months, it is likely that there is pathophysiological progression in the brain, but that more time is needed for the disease to reach the dementia stage. Another possibility might be related to cognitive reserve [27–29]. The cognitive reserve hypothesis suggests that at a particular level of AD pathology, different individuals manifest different levels of clinical symptoms of dementia. It has been shown that highly educated individuals are less likely to manifest clinical symptoms of dementia compared to less-educated individuals [27–29]. However, in our study, there was no difference in education levels between the false positive and true positive groups at 12 months, which argues against this explanation.

Furthermore, our study showed that features which are highly sensitive for short follow-ups – for instance disease severity metric (CDR-SB) - can lose importance during longer follow-ups, while some other features may show low sensitivity to early stage changes and gain importance in longer follow-ups like the entorhinal cortex SNIPE grading and RAVLT score. Furthermore, certain baseline cognitive scores, like RAVLT, are better than others for longer follow-ups. This confirms a previous study that showed that verbal memory measures and language tests have high predictive value for progression to dementia during the MCI period [14, 23]. That said, a simple bedside test like the MMSE still has reasonable accuracy to predict conversion.

For mid-range time periods, the combined MRI and cognitive biomarker model closely predicts the follow-up clinical diagnosis. Our AUC reaches 0.92 for five-year follow up, which, to the best of our knowledge, is the highest AUC reported for five-year follow-up for any predictive method in AD.

This study is not without limitations. While we used the large publicly available multi-site ADNI dataset, the longest follow-up periods would benefit from a larger subject pool as the number of subjects was highly constrained by the availability of diagnosis, study data and availability of MR image at baseline. Since the dataset size was limited in longer follow-ups, we decided to use LOO cross-validation. While the method keeps the test and training sets separate, a less data conservative variant, such as 10-fold cross-validation, could be used in case of availability of larger datasets.

In addition, ADNI includes patients only with amnestic-MCI, therefore we cannot determine the accuracy of our model for Alzheimer’s disease beyond the typical clinical presentation. A more heterogeneous population, more representative of the clinical population with prospective follow-up would be needed to evaluate these tools before they could be used in the clinic. Finally, we do not compare our results to those that use molecular imaging such as amyloid-PET [30]. In future work, we will combine amyloid imaging with our MRI and cognitive features.

## Conclusion

We have demonstrated that combining MRI features with cognitive test results from a baseline visit can be used to predict the onset of dementia in a large cohort over periods up to 9 years, but with maximal accuracy up to five years. The two year FINGER trial [6] demonstrated significant benefit of combined diet, exercise and cognitive training interventions to prevent cognitive decline in cognitively at risk adults. Early detection (e.g., 5-10y) is key to intervene to prevent irreversible neurodegeneration and the onset of cognitive decline. In addition, the uncertainty of future progression is a major source of anxiety for patients, therefore any tool that increase the accuracy of course prediction would be of great benefit. With additional validation, this tool could be useful in the clinic for better patient management and for cohort enrichment in clinical trials of new treatments for AD.

## Acknowledgment

This work was supported by grants from the Canadian Institutes of Health Research (MOP-111169), les Fonds de Research Santé Quebec Pfizer Innovation fund, and an NSERC CREATE grant (4140438 - 2012). We would like to acknowledge funding from the Famille Louise & André Charron.

Data collection and sharing for this project was funded by the Alzheimer’s Disease Neuroimaging Initiative (ADNI) (National Institutes of Health Grant U01 AG024904) and DOD ADNI (Department of Defense award number W81XWH-12-2-0012). ADNI is funded by the National Institute on Aging, the National Institute of Biomedical Imaging and Bioengineering, and through generous contributions from the following: AbbVie; Alzheimer’s Association; Alzheimer’s Drug Discovery Foundation; Araclon Biotech; BioClinica, Inc.; Biogen; Bristol-Myers Squibb Company; CereSpir, Inc.; Cogstate; Eisai, Inc.; Elan Pharmaceuticals, Inc.; Eli Lilly and Company; EuroImmun; F. Hoffmann-La Roche Ltd and its affiliated company Genentech, Inc.; Fujirebio; GE Healthcare; IXICO Ltd.; Janssen Alzheimer Immunotherapy Research & Development, LLC.; Johnson & Johnson Pharmaceutical Research & Development, LLC.; Lumosity; Lundbeck; Merck & Co., Inc.; Meso Scale Diagnostics, LLC.; NeuroRx Research; Neurotrack Technologies; Novartis Pharmaceuticals Corporation; Pfizer, Inc.; Piramal Imaging; Servier; Takeda Pharmaceutical Company; and Transition Therapeutics. The Canadian Institutes of Health Research is providing funds to support ADNI clinical sites in Canada. Private sector contributions are facilitated by the Foundation for the National Institutes of Health (www.fnih.org). The grantee organization is the Northern California Institute for Research and Education, and the study is coordinated by the Alzheimer’s Therapeutic Research Institute at the University of Southern California. ADNI data are disseminated by the Laboratory for Neuroimaging at the University of Southern California.

Dr. Ducharme reports no conflicts of interest related to this study. Dr. Ducharme receives salary funding from the *Fonds de Recherche du Québec-Santé*.

Dr. Collins reports no conflicts of interest related to this study. Dr. Collins provides training and consulting to NeuroRx and is co-founder of True Positive Medical Devices.

## Supplementary Materials

### Feature importance Metric

The measure is developed based on underlying assumption of Naïve Bayes which is the features independence given each class (Equation 1). Therefore, feature importance can be defined as the separability of two Gaussian distributions (Equation 2).

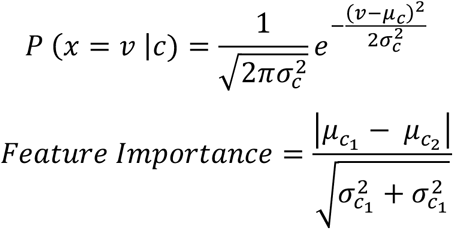

which is similar to an effect size.

## Footnotes

1 Data used in preparation of this article were obtained from the Alzheimer’s Disease Neuroimaging Initiative (ADNI) database (adni.loni.usc.edu). As such, the investigators within the ADNI contributed to the design and implementation of ADNI and/or provided data but did not participate in analysis or writing of this report. A complete listing of ADNI investigators can be found at: http://adni.loni.usc.edu/wp-ontent/uploads/how_to_apply/ADNI_Acknowledgement_List.pdf

